## Supplementary Information for "The Notch1 intracellular domain orchestrates mechanotransduction of fluid shear stress"

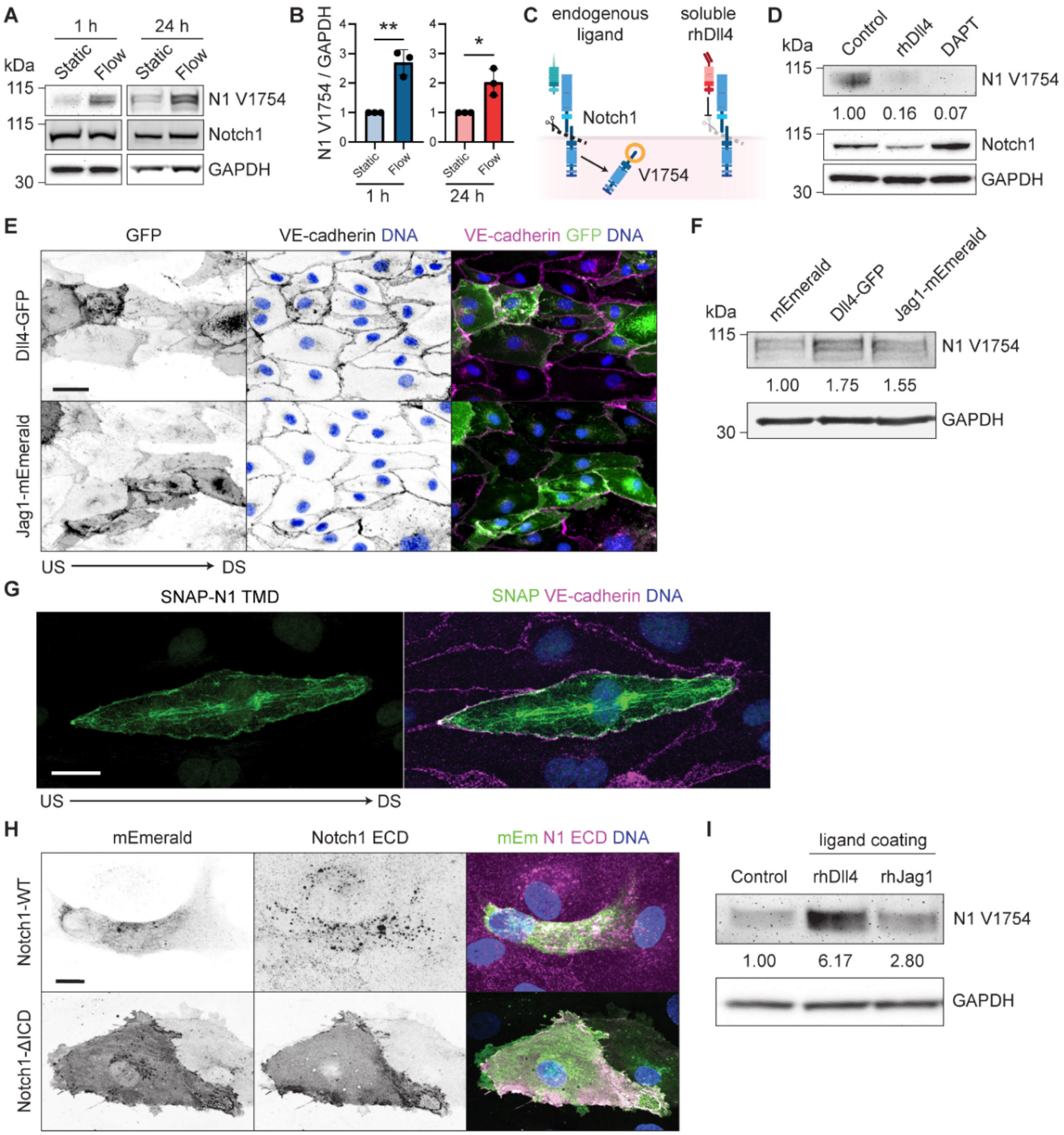

A
1 h
24 h
kDa
Static
Flow
Static
Flow
115
N1 V1754
115
Notch1
30
GAPDH
B
N1 V1754 / GAPDH
**
*
Static
Flow
1 h
24 h
C
endogenous ligand
soluble rhDll4
Notch1
V1754
D
kDa
Control
rhDll4
DAPT
115
N1 V1754
1.00
0.16
0.07
115
Notch1
30
GAPDH
E
GFP
VE-cadherin DNA
VE-cadherin GFP DNA
Dll4-GFP
Jag1-mEmerald
US
DS
F
kDa
mEmerald
Dll4-GFP
Jag1-mEmerald
115
N1 V1754
1.00
1.75
1.55
30
GAPDH
G
SNAP-N1 TMD
SNAP VE-cadherin DNA
US
DS
H
mEmerald
Notch1 ECD
mEm N1 ECD DNA
Notch1-WT
Notch1-ΔICD
I
ligand coating
Control
rhDll4
rhJag1
N1 V1754
1.00
6.17
2.80
GAPDH

**Supplementary Figure 1**

**(A)** Western blot of hdBEC lysates cultured static or following 1 or 24 h of laminar flow. **(B)** Quantification of normalized Western blot intensity of Notch1 V1754. **(C)** Schematic of Notch1-ligand engagement and subsequent proteolytic cleavage, exposing ICD residue V1754, compared to engagement with soluble rhDll4 which blocks ligand binding and subsequent proteolytic cleavage of Notch1. **(D)** Western blot of hdBEC lysates treated with 0.5 ug/mL of rhDll4, 10 μM DAPT, or vehicle control for 3 hours. **(E)** Fluorescence micrographs of hdBEC monolayers expressing Dll4-GFP or Jag1-mEmerald (green). Scale bar, 50 μm. **(F)** Western blot of hdBEC lysates expressing Dll4-GFP, Jag1-mEmerald, or an mEmerald control. **(G)** Fluorescence micrographs of hdBEC monolayer under flow conditions expressing SNAP-N1 TMD (green). Scale bar, 25 μm. **(H)** Fluorescence micrographs of unpermeabilized Notch1-WT and Notch1-ΔICD expressing hdBECs (green) immunostained for construct surface expression using Notch1 ECD antibody (magenta). Scale bar, 25 μm. **(I)** Western blot of hdBEC lysates plated on control, rhDll4-coated, or rhJag1-coated dishes.

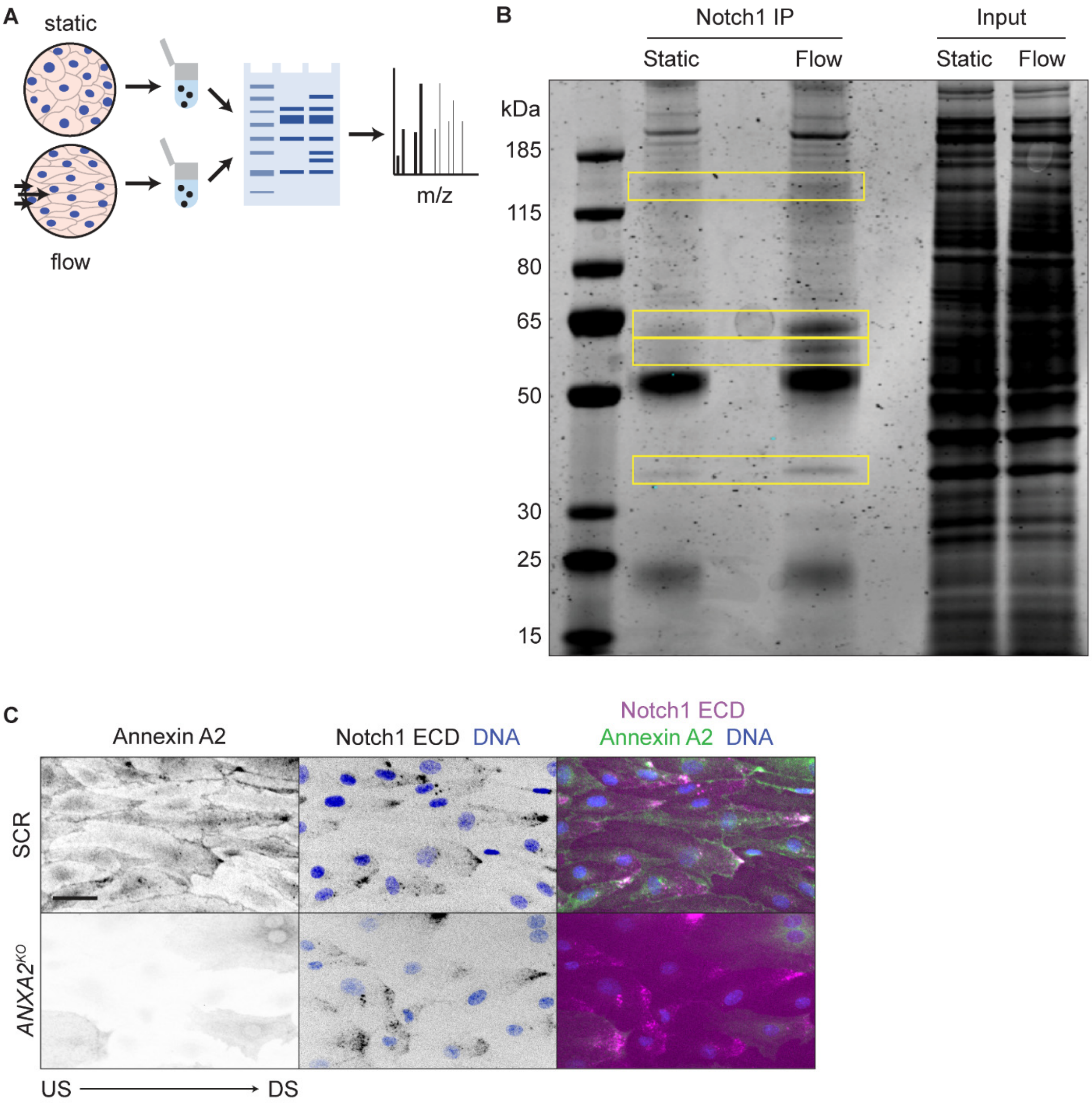
A
static
flow
m/z
B
Notch1 IP
Input
Static
Flow
Static
Flow
kDa
185
115
80
65
50
30
25
15
C
Annexin A2
Notch1 ECD DNA
Notch1 ECD
Annexin A2 DNA
SCR
ANXA2KO
US
DS

**Supplementary Figure 2**

**(A)** Schematic of mass spectrometry workflow to identify flow-dependent Notch1 ICD protein-protein interactions. **(B)** Coomassie stained SDS-PAGE gel of Notch1 co-immunoprecipitation from hdBECs lysates under static and flow conditions. Yellow boxes denote bands analyzed via mass spectrometry. **(C)** Fluorescence micrographs of SCR versus *ANXA2*$^{KO}$ cells under flow conditions. Scale bar, 25 μm.

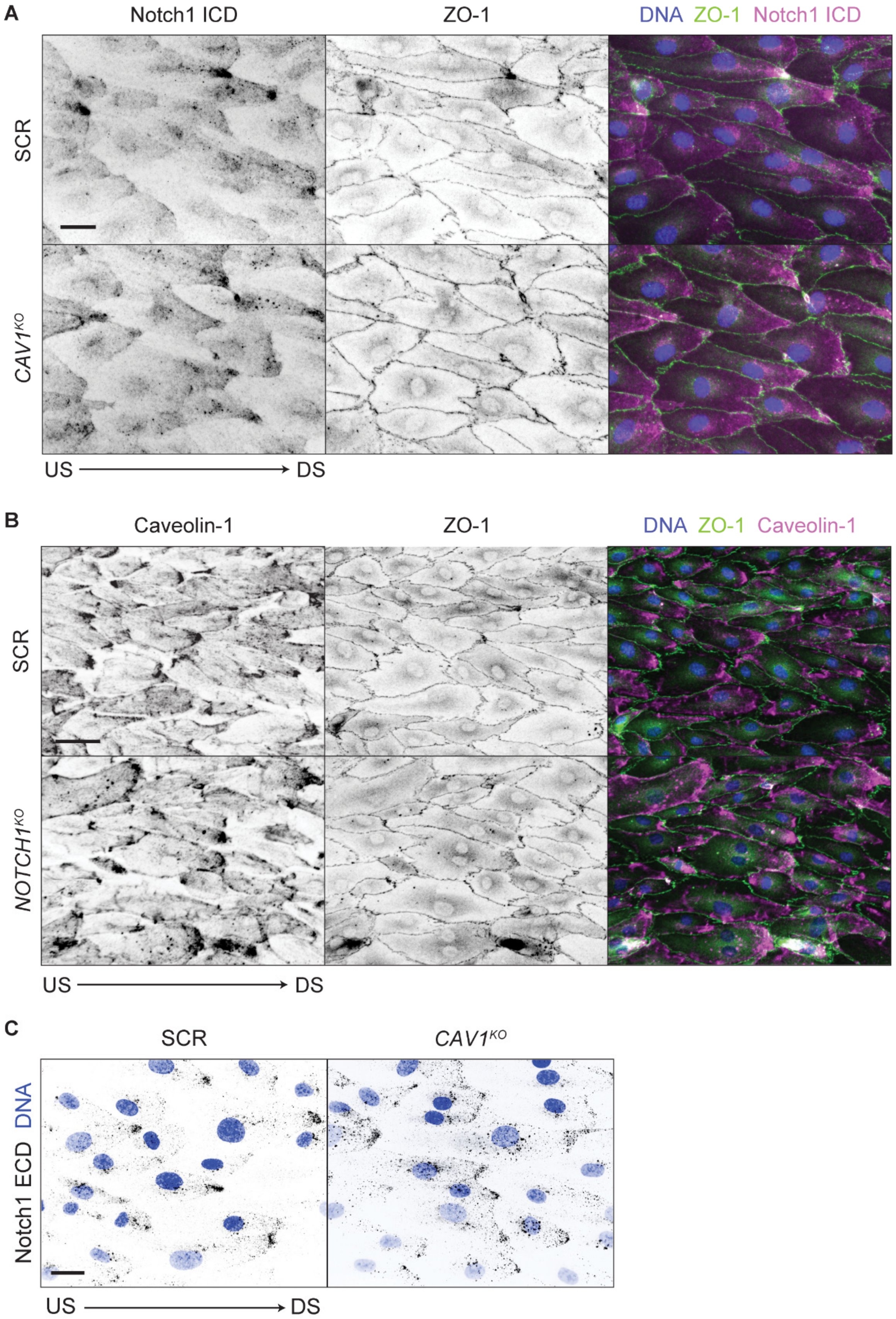
A
Notch1 ICD
ZO-1
DNA ZO-1 Notch1 ICD
SCR
CAV1KO
US DS
B
Caveolin-1
ZO-1
DNA ZO-1 Caveolin-1
SCR
NOTCH1KO
US DS
C
SCR
CAV1KO
Notch1 ECD DNA
US DS

**Supplementary Figure 3**

**(A)** Fluorescence micrographs of SCR versus *CAV1*$^{KO}$ cells under flow immunostained for ZO-1 (green), Notch1 ICD (magenta) and DNA (blue). Scale bar, 25 µm. **(B)** Fluorescence micrographs of SCR versus *NOTCH1*$^{KO}$ cells under flow immunostained for ZO-1 (green), caveolin-1 (magenta) and DNA (blue). Scale bar, 25 µm. **(C)** Fluorescence micrographs of internalized Notch1 in SCR and *CAV1*$^{KO}$ cells under flow visualized with pulse-chase labelling. Scale bar, 25 µm.

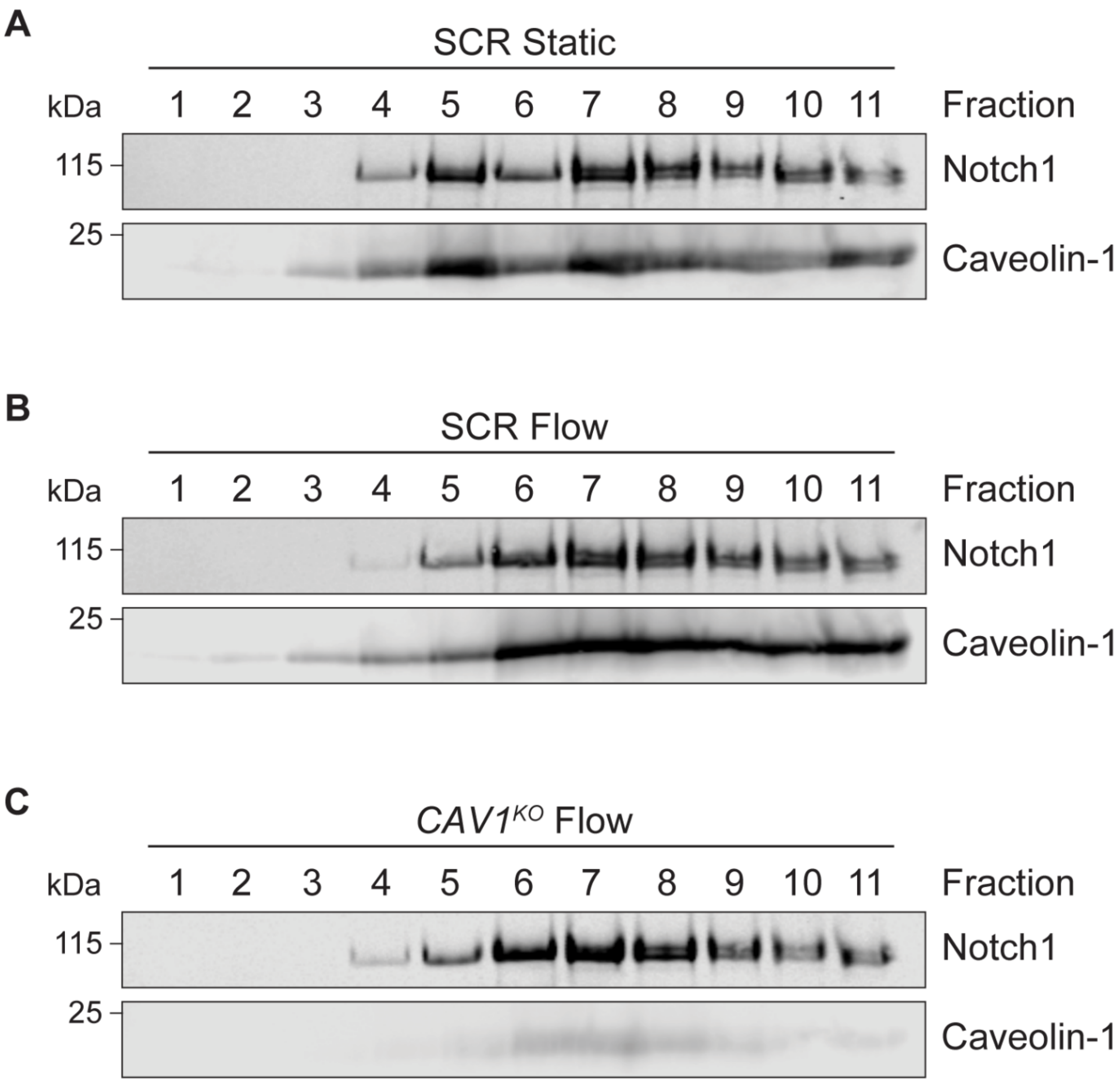
A
SCR Static
kDa
1 2 3 4 5 6 7 8 9 10 11
Fraction
115
Notch1
25
Caveolin-1
B
SCR Flow
kDa
1 2 3 4 5 6 7 8 9 10 11
Fraction
115
Notch1
25
Caveolin-1
C
CAV1^KO Flow
kDa
1 2 3 4 5 6 7 8 9 10 11
Fraction
115
Notch1
25
Caveolin-1

**Supplementary Figure 4**

**(A)** Western blots of membrane fractions isolated from SCR hdBEC lysates under static culture immunoblotted for Notch1 and caveolin-1. **(B)** Western blots of membrane fractions isolated from SCR hdBEC lysates under flow immunoblotted for Notch1 and caveolin-1. **(C)** Western blots of membrane fractions isolated from $CAV1^{KO}$ hdBEC lysates under flow immunoblotted for Notch1 and caveolin-1.

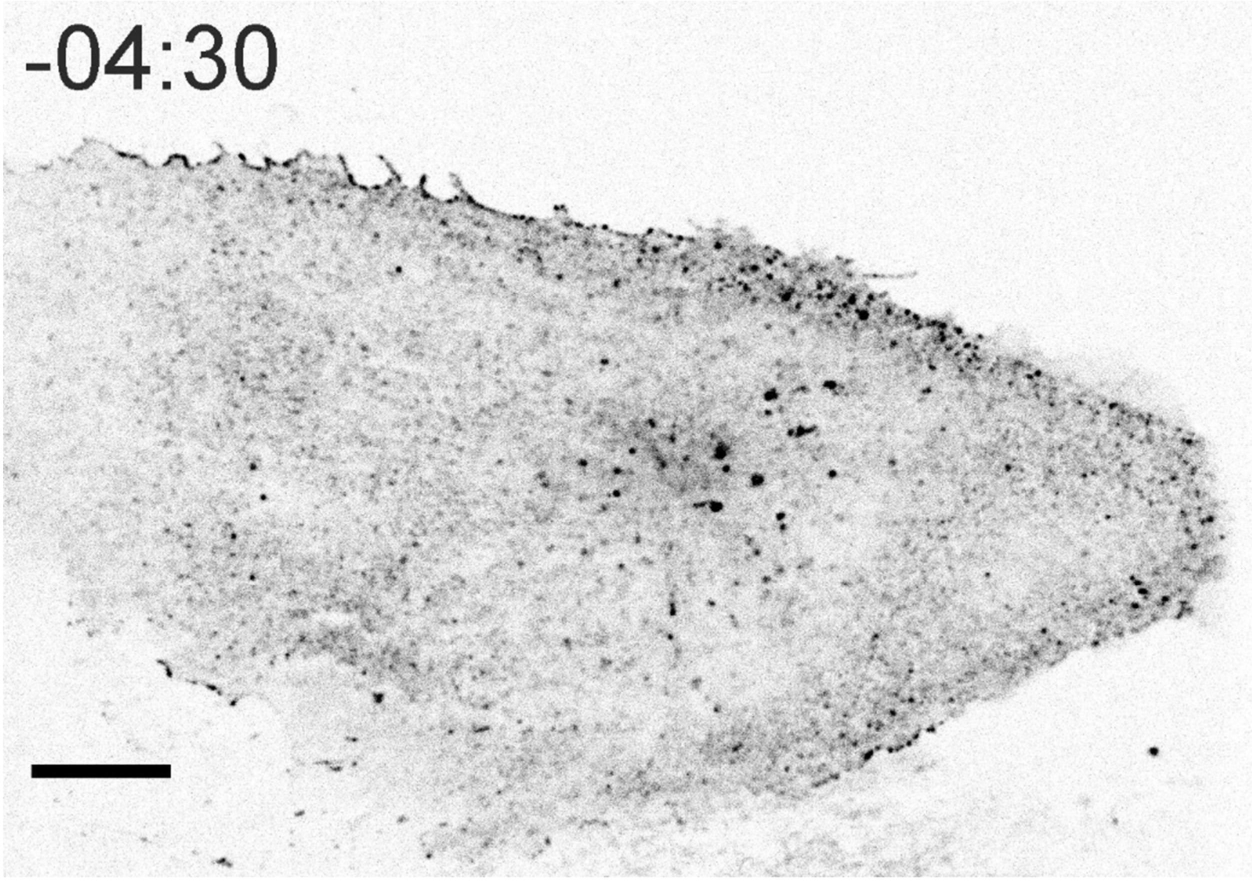


**Supplementary Video 1: Live-cell imaging of hdBEC expressing Notch1-GFP within monolayer under flow**. Video begins under static conditions and flow is initiated at 00:00. Direction of flow is indicated by arrow. Scale bar, 10 μm. Time scale, minute:second, displayed at 5 fps.

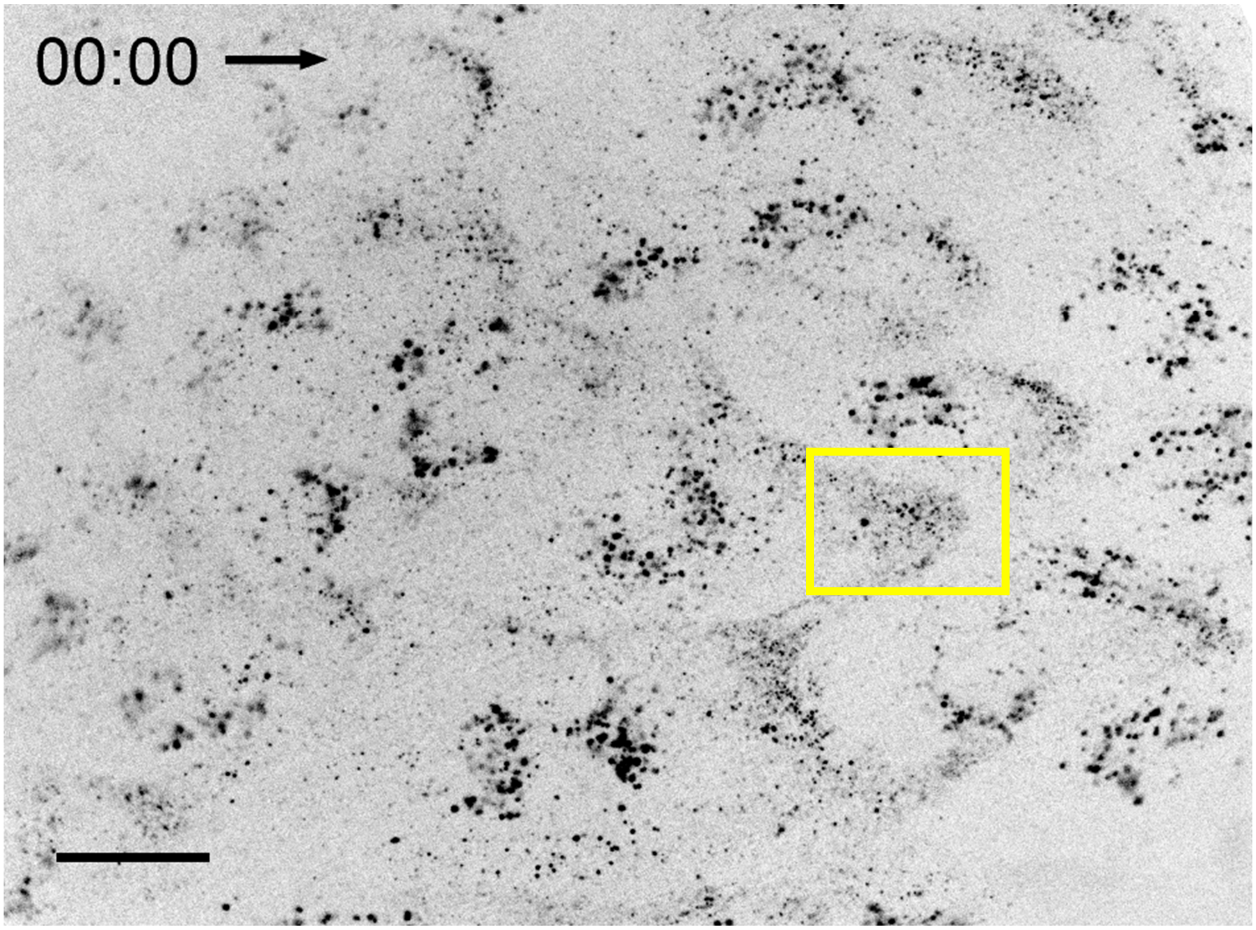


**Supplementary Video 2: Live-cell imaging of antibody-labelled Notch1 within hdBEC monolayer under flow**. Direction of flow is indicated by arrow. Scale bar, 25 µm. Scale bar inset, 5 µm. Time scale, minute:second, displayed at 5 fps.

**Fig. 1A**

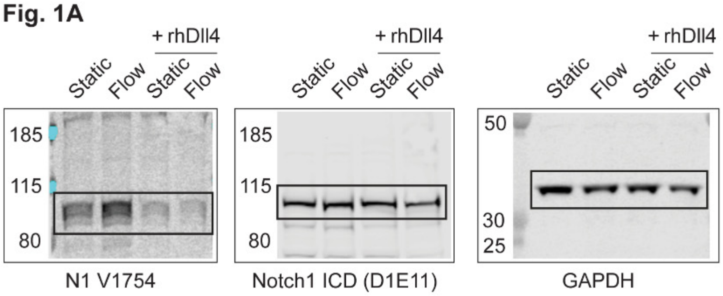


**Fig. 1C**

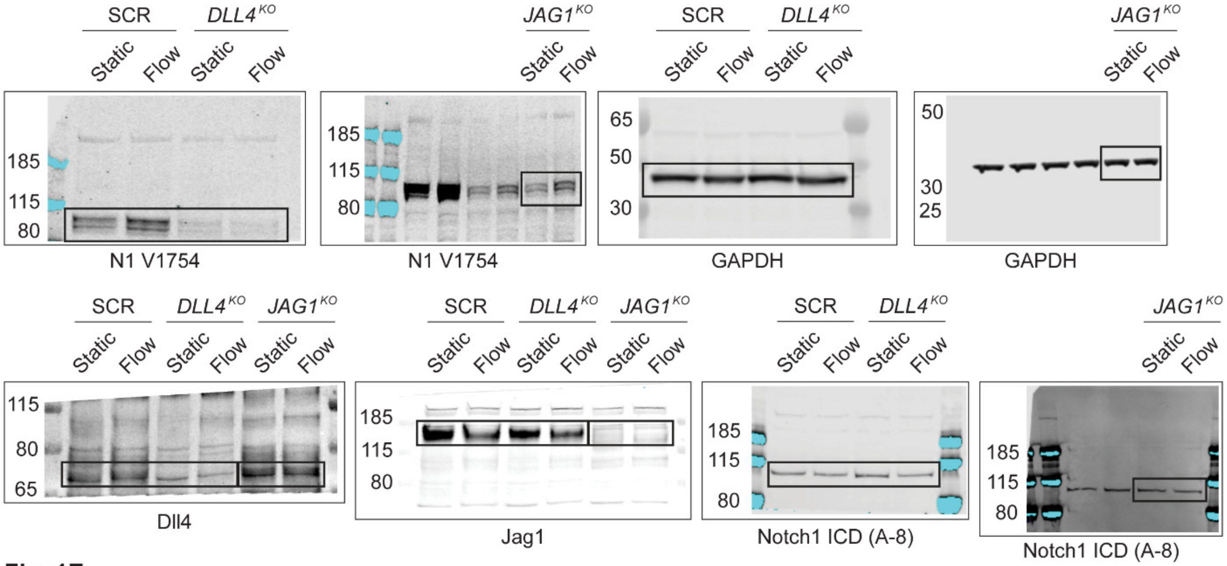


**Fig. 1E**

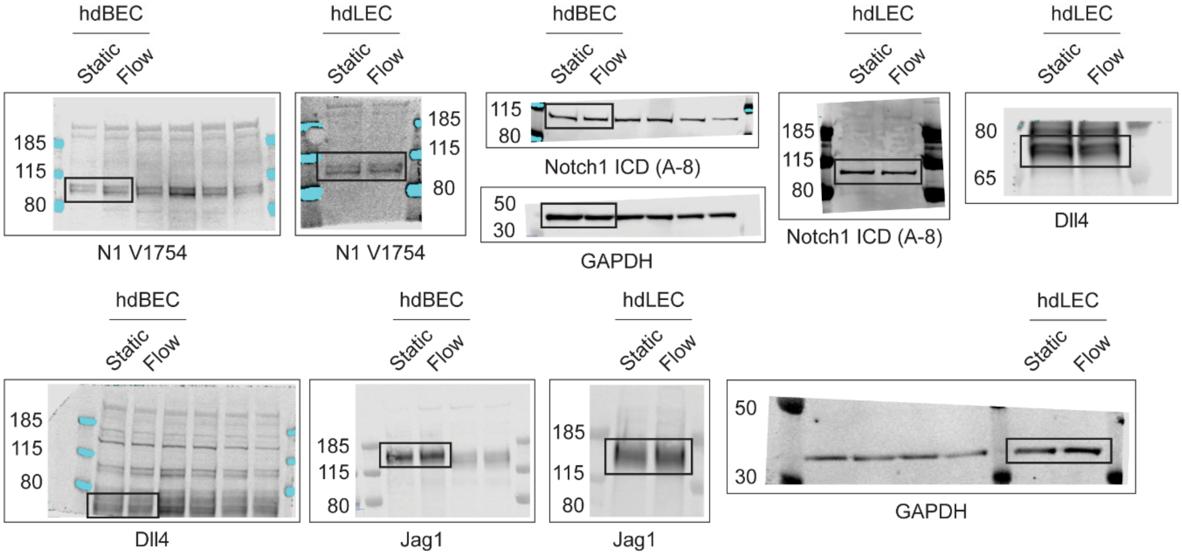

**Supplmentary Fig. 1A**

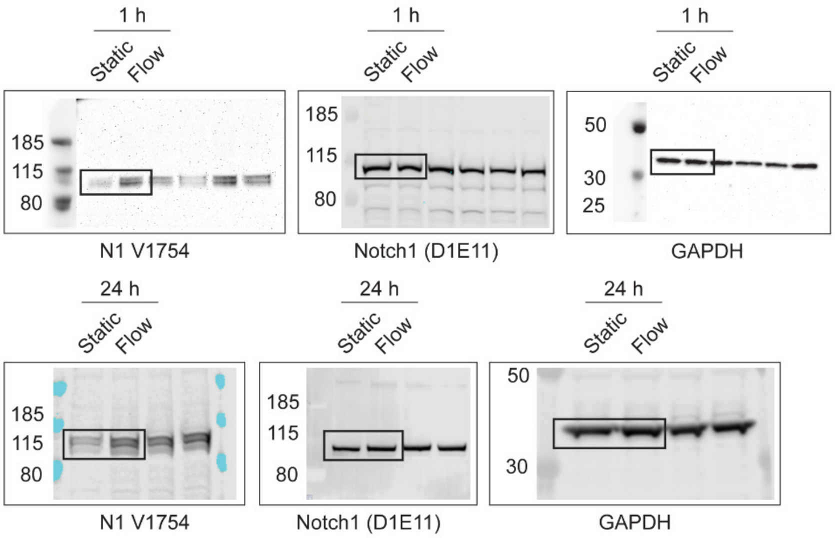


**Supplmentary Fig. 1D**

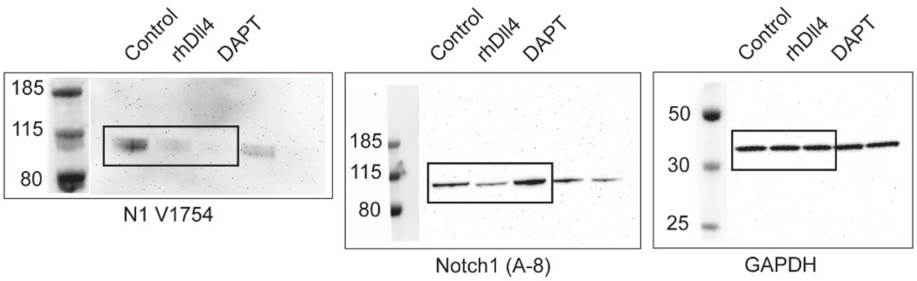


**Supplmentary Fig. 1F**

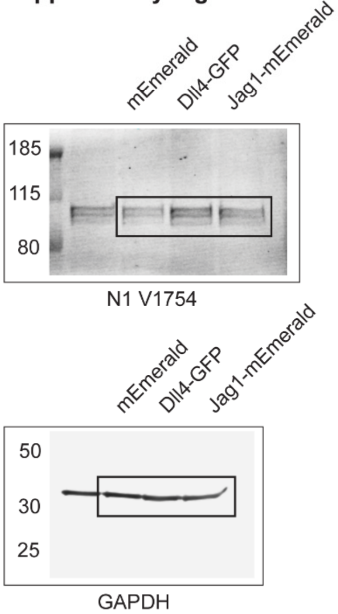


**Supplmentary Fig. 1I**

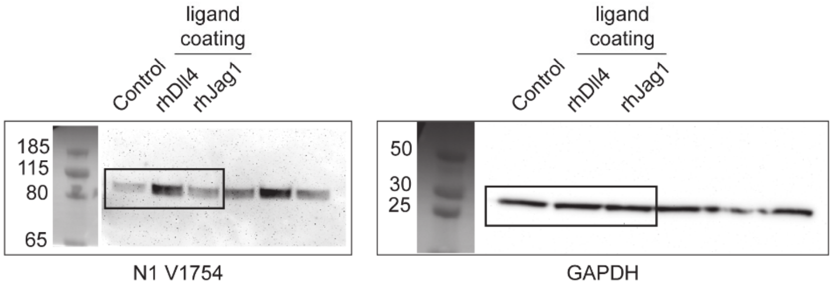

**Fig. 3B**

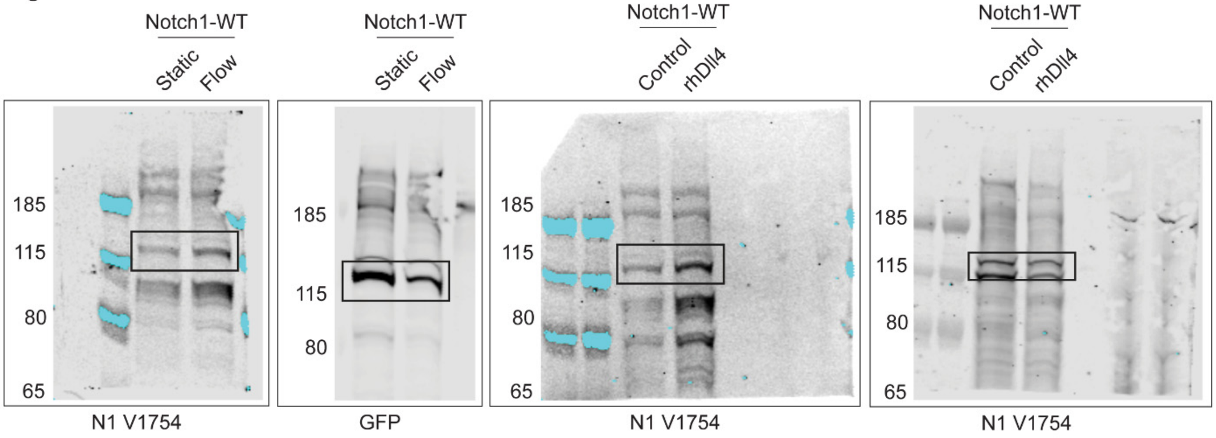


**Fig. 3C**

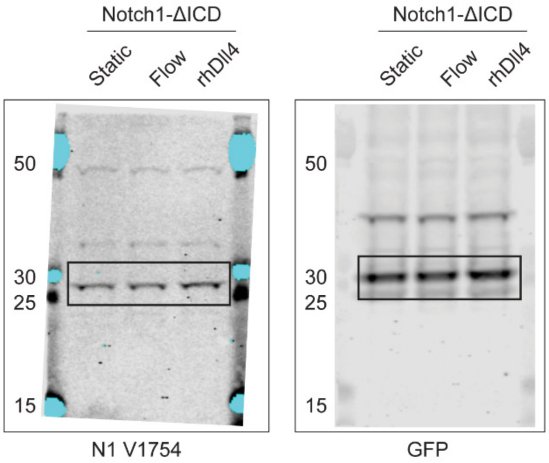


**Fig. 3D**

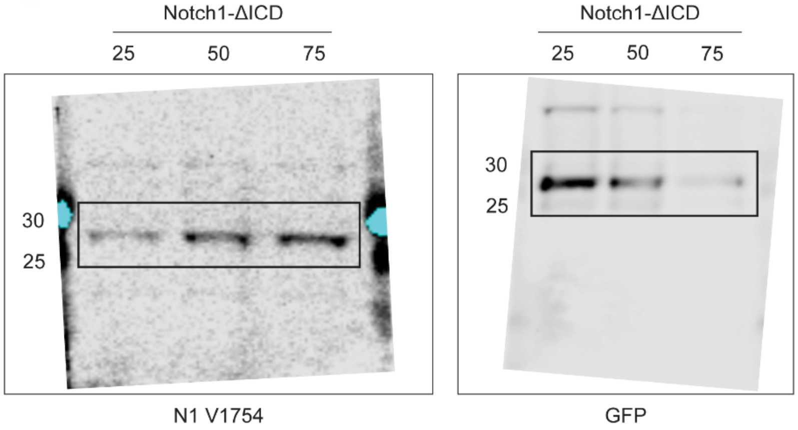

**Fig. 4B**

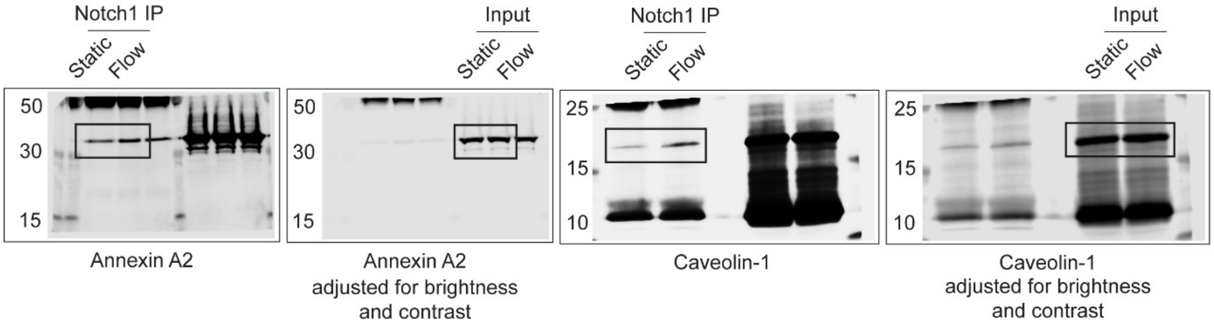


**Fig. 4C**

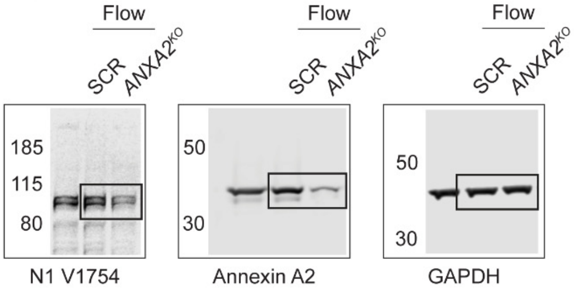


**Fig. 4J**

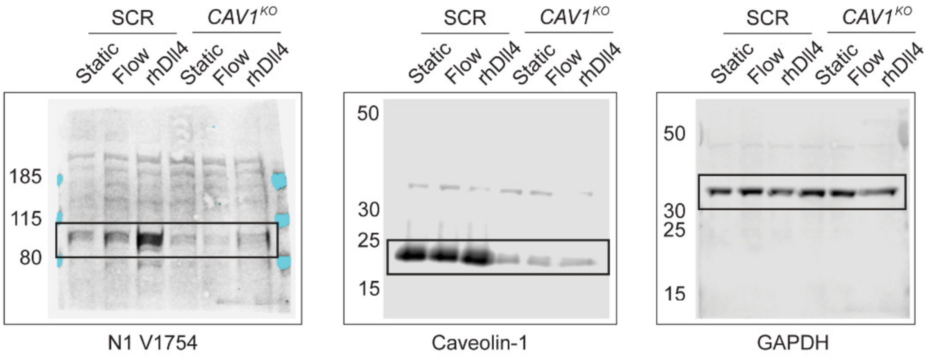


**Fig. 4N**

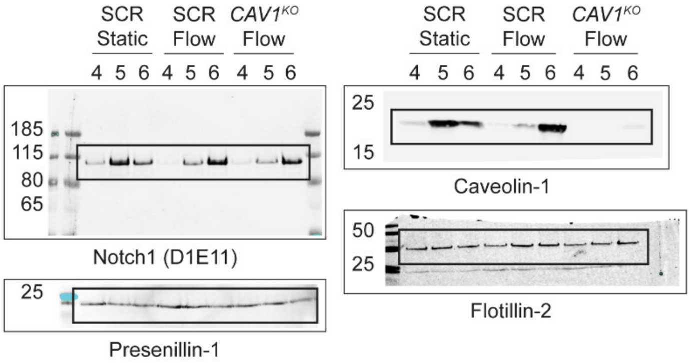

**Supplmentary Fig. 4A**

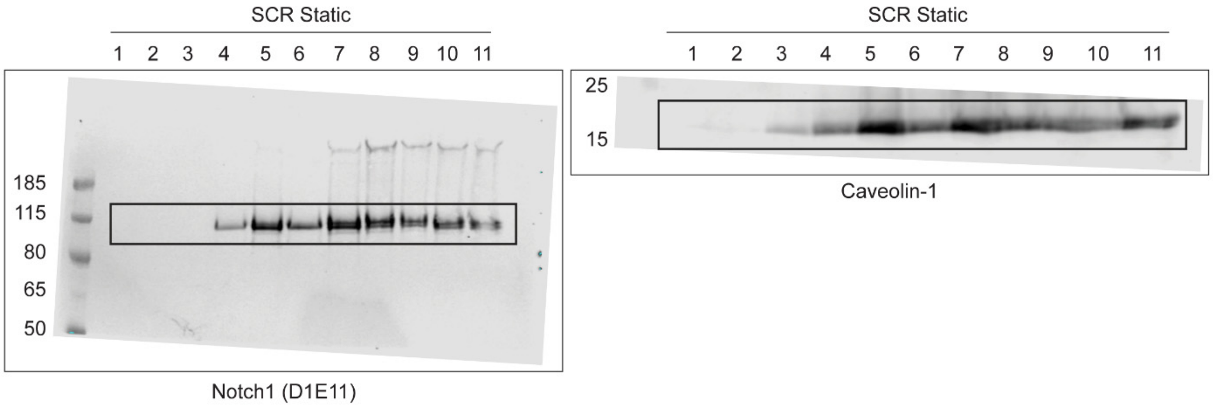


**Supplmentary Fig. 4B**

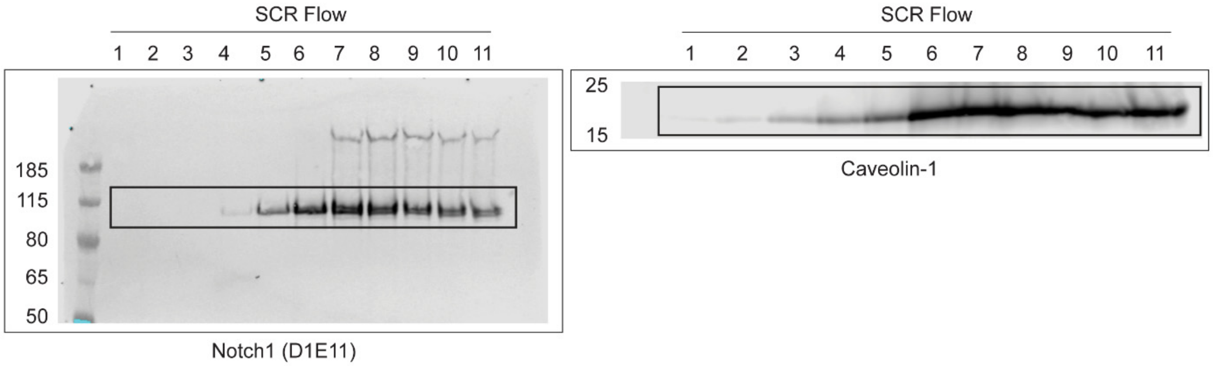


**Supplmentary Fig. 4C**

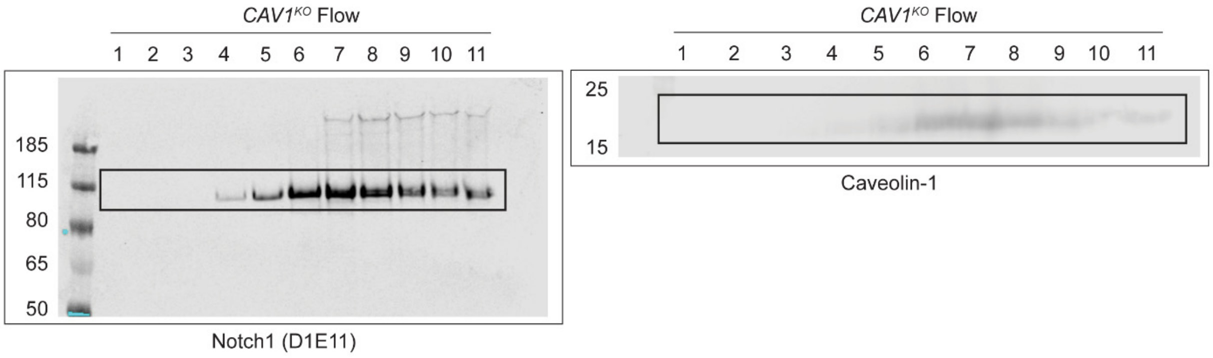
